# Simultaneous Overexpression of Three Enzymes of Chloroplast Metabolism Fails to Improve CO_2_ Assimilation or Biomass

**DOI:** 10.1101/2024.12.11.627974

**Authors:** Pauline Lemonnier, Shellie Wall, Hussein Gherli, Beatriz Moreno-Garcia, Chidi Afamefule, Tracy Lawson, Christine A. Raines, Patricia E. Lopez-Calcagno

## Abstract

Ensuring an adequate food supply amidst a growing global population and climate change challenges, necessitates innovative strategies to enhance crop productivity. Previous studies have demonstrated that the simultaneous stimulation of different photosynthesis-related processes can increase the rate of photosynthetic carbon assimilation and plant biomass. This study evaluates an approach based on modelling aimed at simultaneously increasing photosynthetic and sink capacities in *Nicotiana tabacum* by overexpressing three key enzymes: Sedoheptulose-1,7-bisphosphatase (SBPase), Fructose-1,6-bisphosphate aldolase (FBP Aldolase), and ADP-glucose pyrophosphorylase (AGPase). Our results showed that this strategy does not significantly improve growth or carbon assimilation in *Nicotiana tabacum* under the tested conditions. This suggests that while the model informing our work offers a valuable framework, its application may require adjustments based on species and environmental conditions. Future research should explore these genetic modifications in species with larger sink capacities and under a range of growth conditions to fully realize the potential of photosynthetic optimization.

## INTRODUCTION

As our population continues to increase, ensuring an adequate food supply becomes increasingly challenging. To secure future food supplies without increasing current agricultural lands, in a world affected by climate change, the continued development of innovative strategies to enhance crop productivity is needed. Central to this challenge are the processes of photosynthesis and the intricate source-sink relations that regulate carbon allocation within plants. Understanding how we can optimize these processes has the potential to lead to significant improvements in agricultural output.

Over the last two decades, a number of empirical studies have shown consistently how increasing the investment in some Calvin-Benson-Bassham (CBB) cycle enzymes (by overexpression of these) is able to increase photosynthetic productivity. The first efforts focused on manipulation of single enzymes, a good example of which is the overexpression of sedoheptulose-1,7-bisphosphatase (SBPase) in four different species, *Arabidopsis thaliana (Simkin et al., 2017)*, tobacco (Lefebvre et al., 2005; Rosenthal et al., 2011; Simkin et al., 2015), tomato (Ding et al., 2016) and wheat (Driever et al., 2017), which consistently showed increases in both photosynthetic carbon assimilation and yield. Other targets, like the introduction of the cyanobacterial fructose-1,6-bisphosphatases/sedoheptulose-1,7-bisphosphatase (cyFBP/SBPase) enzyme (Miyagawa et al., 2001; Tamoi et al., 2006; Ichikawa et al., 2010; Köhler et al., 2017), and the overexpression of fructose-1,6-bisphosphate aldolase (FBP Aldolase) (Uematsu et al., 2012; Simkin et al., 2015; Simkin et al., 2017; Cai et al., 2022) have also shown increases in productivity. However, more recently, work focused on the simultaneous stimulation of multiple enzymes has shown increased success in photosynthesis stimulation. Simkin *et al*. (Simkin et al., 2015; Simkin et al., 2017), showed how the combined overexpression of SBPase and FBP Aldolase in *Arabidopsis thaliana* and tobacco led to further increases in productivity than the single overexpression of either of these enzymes. Furthermore, these studies, and others have also shown a positive effect in photosynthetic carbon assimilation and yield by not only stimulating the CBB cycle, but when combining this with targets in other photosynthetic processes like photorespiration, by increasing the levels of the H-protein of the glycine cleavage system (Simkin et al., 2017); electron transport chain, by introducing cytochrome *c*_6_ in higher plants (Lopez-Calcagno et al., 2020), or by combination with the putative inorganic carbon transporter B (Simkin et al., 2015) (ictB).

In agreement with these empirical evidence and using a modelling approach, Zhu et al.’s (2007) proposed that “*the distribution of resources between enzymes of photosynthetic carbon metabolism might be assumed to have been optimized by natural selection. However, natural selection for survival and fecundity does not necessarily select for maximal photosynthetic productivity*”. Taking this into account, as well as the rapid changes to atmospheric CO_2_ concentrations which we are currently experiencing, calls for a review of the enzyme balance for maximizing the light-saturated rate of photosynthesis. The authors proposed that currently plants have an overinvestment in photorespiratory pathway (PCOP) enzymes and underinvestment in CBB cycle enzymes like Rubisco, SBPase and FBP Aldolase. Furthermore, it proposed that an increase in sink capacity, such as that obtained by increasing ADP-glucose pyrophosphorylase (AGPase), would also be needed for an increased CO_2_ uptake rate. Specifically, the model included 23 enzymes and proposed that for optimizing photosynthesis at around current atmospheric CO_2_ concentrations, the levels of seven of these enzymes, all belonging to the CBB cycle with the exception of AGPase, would need increasing, while the levels of the 14 other enzymes decreased, two of which belong to the CBB cycle and seven to the PCOP.

In addition to this proposal (Zhu et al., 2007); there is also empirical evidence that the limitation on P_i_ recycling due to conditions where the potential for photosynthesis exceeds the rate of end-product synthesis, like limitations on the capacity for sucrose and starch synthesis (sink), can induce a negative feedback response in net photosynthesis (Paul and Foyer, 2001; Pieters et al., 2001; Paul and Pellny, 2003). Evidence of the role of leaf starch on influencing the capacity for photosynthesis has clearly been shown using starch deficient and starch null Arabidopsis mutants. In these plants the rates of CO_2_ assimilation and O_2_ evolution were proportional to the level of AGPase in the plants; being highest for wildtype, and lowest for the starch null mutants (Sun et al., 1999). This strongly suggested that leaf starch is not only important as a transient reserve whose metabolism supports heterotrophic growth in the dark, or transient sink, but also as an essential end-product in triose-P utilization (Gibson et al., 2011).

With this in mind, we have taken a “minimal approach” to simultaneously increase photosynthetic and sink capacity in *Nicotiana tabacum*. Given the strong empirical evidence that downregulating enzymes of the PCOP can have a negative effect in photosynthesis and plant growth (Bauwe, 2023) and that overexpression of certain enzymes in the CBB cycle can lead to significant increases in photosynthesis and plant growth (Uematsu et al., 2012; Simkin et al., 2015; Simkin et al., 2017; Cai et al., 2022), we have focused on overexpression of the three enzymes modelled to require the largest increases: SBPase, FBP Aldolase and AGPase. This paper covers the main findings of these experiments as well as suggest that species choice will play a crucial role in the success of approaches trying to optimise photosynthesis manipulation with source-sink relationships.

## RESULTS

### Production and selection of tobacco transformants

To explore the impact of simultaneously overexpressing SBPase, FBP Aldolase and AGPase, an overexpression construct for these three genes (Supplemental Figure 1) was used for tobacco transformation. Over 60 independent T0 lines were generated of which 37 had a single T-DNA insert. 10 single T-DNA insert lines were selected at T0 stage based on detectable expression of all three transgenes and T1 seed grown to generate homozygous lines. Nine homozygous lines alongside eight azygous segregants (which have lost the T-DNA of interest and from this point forward were used as control plants) were successfully identified and further characterized. Transgene expression analysis by qPCR in T1 plants confirmed expression of all three transgenes in the nine selected lines, with no expression in selected control lines (Figure 1a). Immunoblot analysis further confirm the identity of these lines with increased levels of the three target proteins in the homozygous lines and no changes in the control lines (Supplemental Figure 2). Lines were further characterized in the T2 generation.

**Figure 1.**
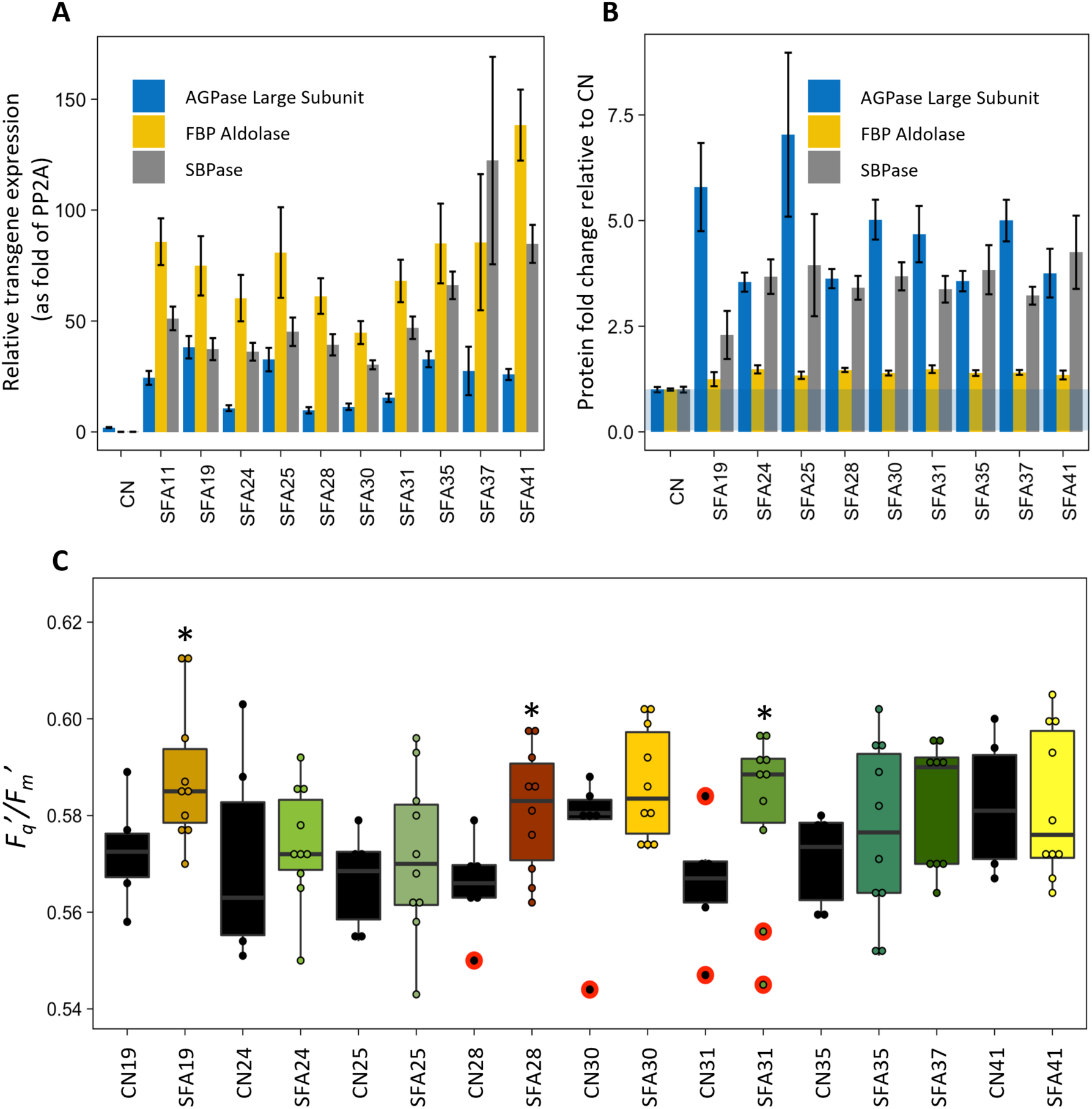
Production and selection of triple overexpressor. **A)** qPCR/transcript abundance in T1 plants. **B)** Protein quantification, relative spectral counts for the target proteins in fully expanded leaves of T2 plants. **C)** *F_q_’/F_m_’* at low O_2_ and 500 μmol m^−2^ s^−1^ of T2 plants 21 to 22 days after sowing. n=6-10. * p<0.05 (linear mixed models accounting for block effect, comparisons done only between each SFA and corresponding CN).

Proteome analysis using tandem mass spectrometry coupled with liquid chromatography (LC-MS/MS) confirmed the consistent accumulation of the three target proteins in T2 homozygous transgenic lines, with no other consistent changes in levels for any other protein. Across the selected lines the abundance of SBPase increased by 2.3- to 4.2-fold, FBP Aldolase by 1.2- to 1.5-fold, and AGPase large subunit by 3.5- to 7-fold compared to the controls (CN) (Figure 1b). It also confirmed that the levels of both AGPase subunits were elevated, suggesting that this approach led to an increase of the functional form of this protein. However, the increase in the AGPase small subunit did not match that of the overexpressed large subunit, only reaching around 2-fold of CN (Supplemental Figure 3). In addition to this, a RNAseq analysis was done in a subset of the lines (SFA19, SFA28 and SFA30) to confirm only the three target genes (SBPase, FBP Aldolase and AGPase) were consistently changed in the lines produced (Supplemental Table 1).

### Photosynthetic characterization: Chlorophyll fluorescence, A/C_i_ response curves, PPFD step response curves and diurnal CO_2_ assimilation

To test if the changes in enzyme levels had a positive effect on photosynthesis chlorophyll fluorescence imaging was used as a quick screen on young plants (21 to 22 days after sowing) grown under control conditions (22°C, 250 μmol m^−2^ s^−1^ and a 16h photoperiod). This analysis was done at low O_2_ and an irradiance of 500 μmol m^−2^ s^−1^ and it showed larger average operating efficiency of PSII photochemistry (*F*_q_’*/F*_m_’) for the transgenics than the grouped controls (CN), with significant increases *F*_q_’*/F*_m_’ in three of the nine lines (Figure 1c).

To further test the impact of increased levels of SBPase, FBP Aldolase and AGPase abundance in carbon fixation, transgenic and control plants were grown in greenhouse conditions (16 h photoperiod, 25 °C - 33 °C day/20 °C night, with natural light supplemented under low light induced by cloud cover with high-pressure sodium light bulbs, giving 380 - 1,000 μmol m^−2^ s^−1^ from the pot level to the top of the plant) and photosynthesis determined using gas exchange on a young developing and a mature leaf of each plant. The data shows that in the young developing leaves, the rate of CO_2_ assimilation (*A*) at different intercellular CO_2_ partial pressures (*C_i_*) and saturating irradiance, was increased compared to CN in in six out of the nine lines measured (Figure 2a and Supplemental Figure 4a). However, in mature leaves, no consistent increase in *A* or change in stomatal conductance (*g*_s_) was evident (Figure 2b and Supplemental Figure 4b). Analysis of the photosynthetic parameters of V*_cmax_*, J*_max_* and TPU on the selected lines in young developing leaves, showed a significantly higher V*_cmax_* in line SFA28 vs CN, and higher average values for all three parameters in four out of the five lines analysed vs CN (Table 1). Contrastingly, in mature leaves, although some lines displayed higher average values than CN, there was no consistent trend and no significant differences for any of the lines in the three parameters analysed (Table 1). This was repeated in two additional experiments with similar results.

**Figure 2.**
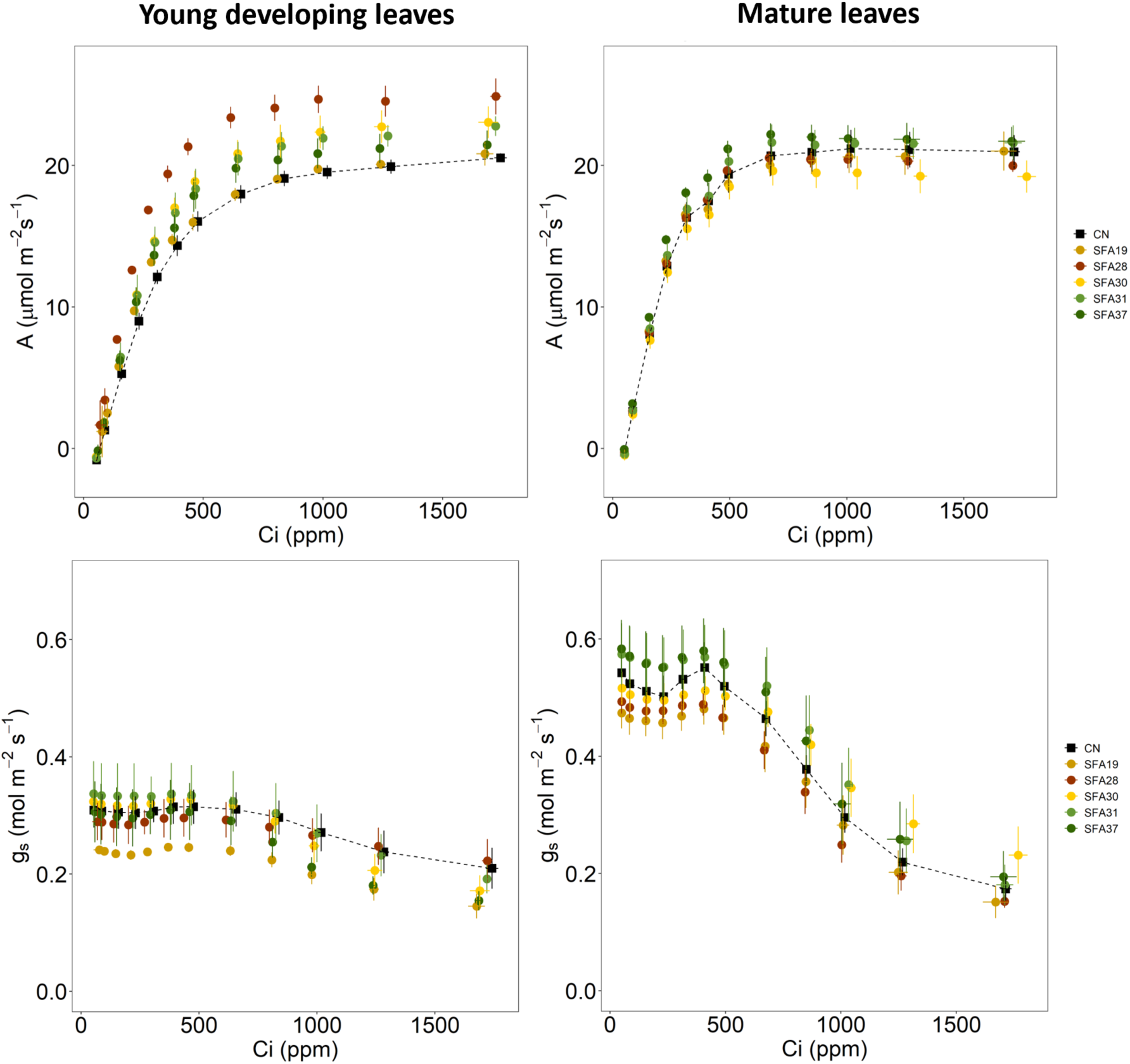
CO_2_ assimilation (*A*) and stomatal conductance (*g_s_*) as a function of increasing intercellular CO_2_ concentrations (Ci) in young developing leaves (left) and mature leaves (right) of transgenic (SFA) and control plants (CN) grown in the glasshouse. Measurements were made at an irradiance of 1500 µmol m^−2^ s^−1^ with a range in leaf temperatures of 26-28 °C. N=3-5.

**Table 1.** Maximum Rubisco carboxylation rate (*V_cmax_*), maximum electron transport rate (*J_max_*) and triose phosphate utilisation (TPU) in young expanding and mature leaves of control (CN) and SFA lines. These parameters are reported at 25 °C and were derived from the *A/C_i_* response curves from Figure 2. Statistical differences obtained from one-way ANOVAs between SFA lines and the control plants are shown in boldface (* P<0.05). Average ± standard error values and number of replicates after removing outliers are shown. NI: Not included

| Genotype | Parameters<br>($\mu\text{mol m}^{-2} \text{s}^{-1}$) | | | | | |
| --- | --- | --- | --- | --- | --- | --- |
|  | Young expanding leaves |  |  | Mature leaves |  |  |
| | $V_{cmax}$ | $J_{max}$ | TPU | $V_{cmax}$ | $J_{max}$ | TPU |
| **CN** | 47.52 ± 2.88 | 93.78 ± 3.20 | 7.37 ± 0.09 | 62.08 ± 4.80 | 101.80 ± 7.77 | 7.42 ± 0.43 |
| **SFA19** | 54.14 ± 1.63 | 91.48 ± 0.85 | NI | 66.16 ± 2.30 | 94.67 ± 0.86 | 6.69 ± 0.07 |
| **SFA28** | **68.59 ± 2.71\*** | 108.51 ± 8.14 | 7.91 ± 0.61 | 62.51 ± 1.53 | 98.65 ± 1.56 | 6.92 ± 0.16 |
| **SFA30** | 59.74 ± 4.30 | 108.02 ± 6.56 | 8.05 ± 0.50 | 60.81 ± 2.70 | 96.59 ± 4.53 | 6.73 ± 0.37 |
| **SFA31** | 59.13 ± 7.27 | 106.07 ± 4.21 | 8.13 ± 0.01 | 66.03 ± 4.97 | 106.04 ± 7.73 | 7.42 ± 0.43 |
| **SFA37** | 54.23 ± 4.31 | 98.85 ± 5.64 | 7.41 ± 0.34 | 70.92 ± 2.36 | 109.33 ± 2.61 | 7.15 ± 0.08 |
| **N** | 3-5 | 2-4 | 2-4 | 3-5 | 3-5 | 3-4 |

To further explore whether mature leaves in the transgenics lines might have changes in *A* during non-steady-state conditions, lines SFA19, SFA28, SFA30, SFA31 and SFA37 were subjected to an increase in photosynthetic photon flux density (PPFD). No significant changes in the speed or amplitude of *A* or *g*_s_ induction were found, when plants were moved from low to high light (Supplemental Figure 5). Additionally, we looked at the integrated diurnal CO_2_ assimilation in matured leaves and assimilation at midday, which also revealed no significant differences (Supplemental Figure 6). While assessing photosynthesis, stomatal impressions were collected to evaluate stomatal density; no significant differences were found (Supplemental Figure 7).

### Carbohydrate measurements

AGPase catalyses the first committed step of starch biosynthesis in higher plants, hence we hypothesized that increasing AGPase capacity by rising the levels of this enzyme would allow any additional carbon fixed to be moved into starch by the removal of the bottlenecks represented by SBPase and FBP Aldolase or negative feedback on photosynthesis by Pi accumulation. Consequently, leaf carbohydrate measurements in mature leaves were done at the end of the day and night. Given the relevance of starch, we first focused on this carbohydrate; Figure 3 shows how line SFA30 displayed significant increases in the starch levels over the CN at the end of the day. However, although it appears to be a trend for starch to be elevated in three of the four lines at the end of the night, no other significant differences with the CN were detected in any of the other lines measured (SFA19, SFA28 and SFA31). In a similar manner, no consistent significant changes were detected in either timepoint in the soluble sugars (sucrose, fructose and glucose) (Supplemental Figure 8).

**Figure 3.**
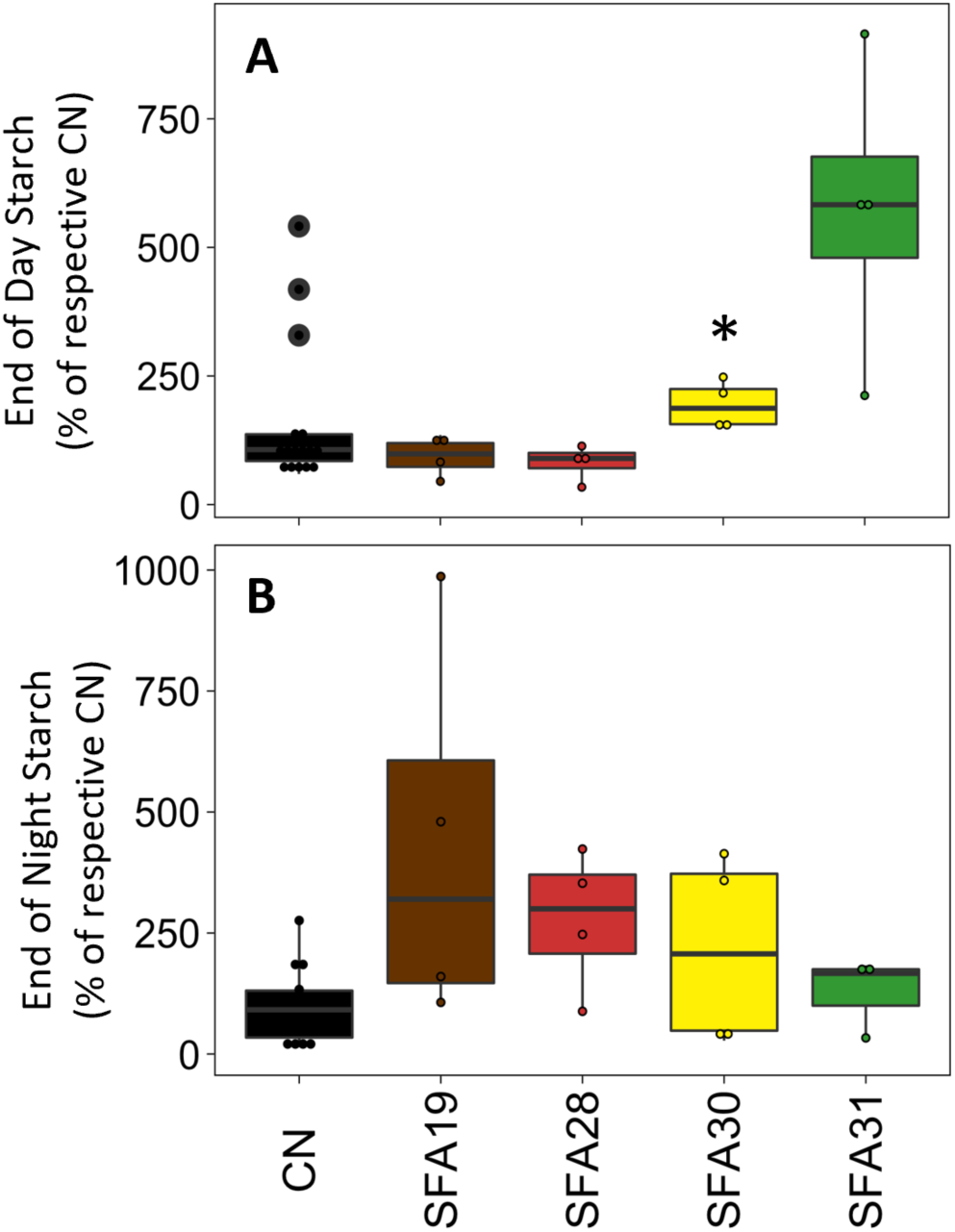
Overexpression of SBPase, FBP Aldolase and AGPase does not consistently change starch profile at the end of the day (A) or end of the night (B). n= 3-4. * p < 0.05 (linear mixed models accounting for block effect, comparisons done only between each SFA and corresponding CN, for visualization of controls separately, see Supplemental figure 8).

### Biomass yield determination

Given that significant increases in *F*_q_’*/F*_m_’ were observed in young plants at light intensities of 500 μmol m^−2^ s^−1^, and consistent increases in *A/C_i_* response curves were only found in young expanding leaves, but no consistent differences in photosynthesis were found later in mature leaves, total leaf area and over ground biomass was measured at three different times in development −25, 46 and 56 days after sowing-for a subset of the transgenic lines (SFA19, SFA28 and SFA30) (Figure 4). The biomass of the full set of lines was also measured at flowering onset, circa 56 days after sowing (Supplemental Figure 9). No impact to total leaf area or over ground biomass was found in any of the timepoints for any of the lines. Additionally, to investigate if any of the potential biomass gains in early development could be allocated the root system, the root biomass was measured 37 days after sowing. No consistent significant differences between the transgenics and CN plants were found (Supplemental Figure 10).

**Figure 4.**
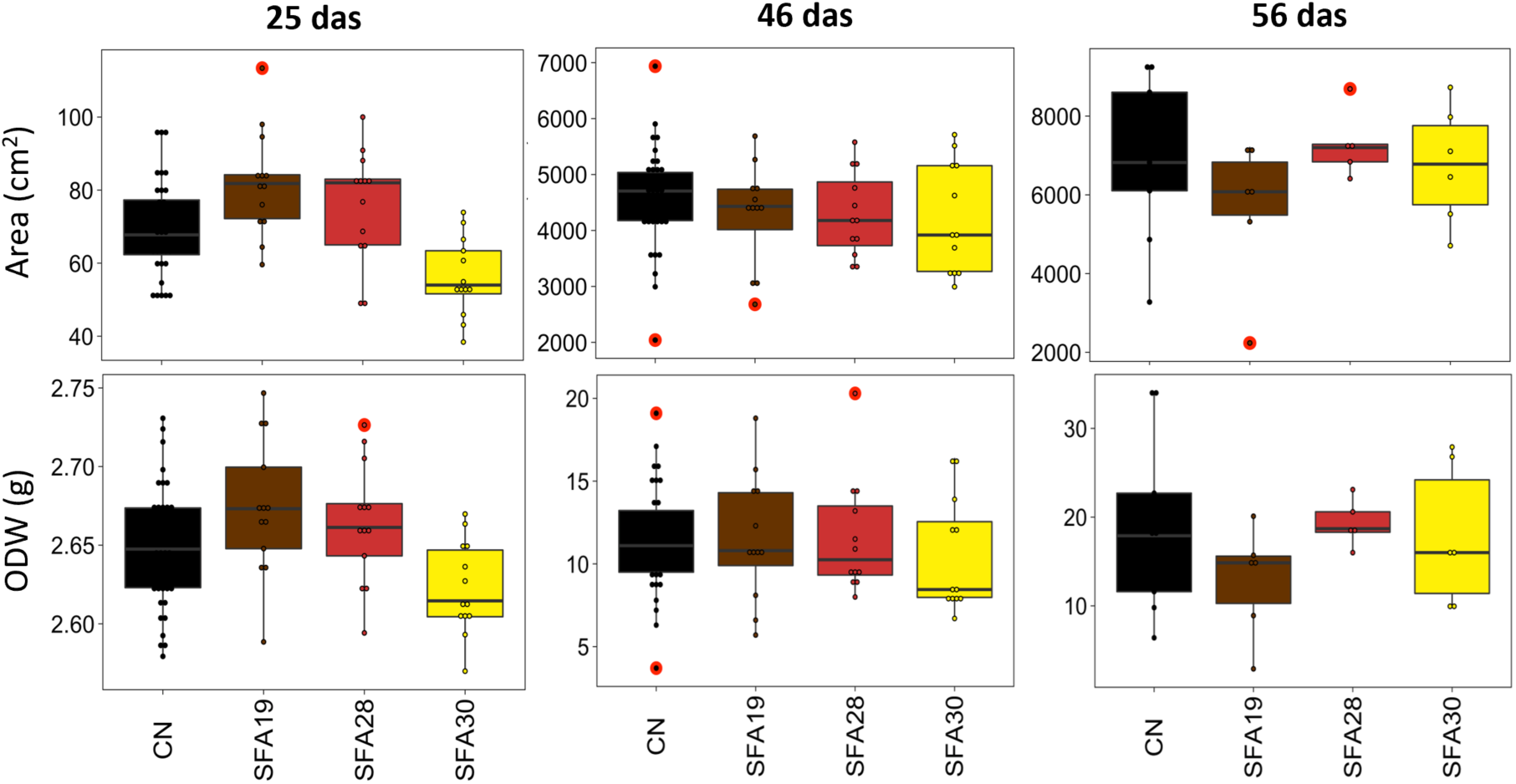
Simultaneous increases in SBPase, FBP Aldolase and AGPase do not affect tobacco biomass in glasshouse-grown plants. At 25 days after sowing (n=13), 46 days after sowing (n= 12) and 56 days after sowing (n= 6 for each SFA line, n=9 for CN). Leaf area (top) and Total over-ground dry weight right (ODW). CN group corresponds to pulled azygous from each line which were not significantly different, statistical analysis compared each overexpressing line to its corresponding CN at 25 and 46 days after sowing. In the 56 das timepoint the CN were not separated by line and hence the comparison was made against the pulled CN group.

## DISCUSSION

The purpose of present work was to assess whether the overexpression of the three enzymes modelled by Zhu *et al*. (2007) to require the largest increases - SBPase, FBP Aldolase and AGPase - would be sufficient to increase photosynthetic carbon assimilation and yield in the model species *Nicotiana tabacum.* Our results clearly show that for this model species and the levels of expression achieved for the target proteins, this approach is not sufficient to significantly affect these parameters. These results, however, shed light on important considerations should this or similar approaches be taken in future to increase photosynthetic carbon assimilation and crop yields.

First, the model was designed to “*partition optimally with respect to maximizing light-saturated photosynthetic rate for a typical C3 leaf*”. However, as in any closed canopy and relevant agricultural setting, not all leaves were continuously light saturated through the life of these plants. Furthermore, under our growth conditions, and specially once moved to the greenhouse, is likely that most the plants were not light saturated the majority of days. However, the steady state measurements of mature leaves at saturating light and dynamic response to saturating light levels (Figure 2 and supplemental Figures 4 and 5) do not support this, failing to display consistent significant increases to *A* at ambient CO_2_ levels and suggesting additional factors at play.

Another key aspect to consider is that the levels of overexpression achieved in these plants or the balance between target proteins might not have been sufficient for the realization of the expected phenotype. For FBP Aldolase the levels of protein obtained ranged between 1.1 and 1.7 fold of control (CN) levels, while Zhu et al.’s (2007) model suggested an increase of over 1.8 fold of this protein. Although not quantified, Simkin et al. (2015), showed large increases in FBP Aldolase protein in transgenic lines overexpressing FBP Aldolase and SBPase plants but with only 1.5 fold in FBP Aldolase enzyme activity. It is possible that larger increases in the level of this enzyme, or the use of alternative FBP Aldolases with higher catalytic activities would be necessary for this approach to yield increases in photosynthetic assimilation. Similarly, it is possible that the increase in active AGPase was not sufficient. The proteomics results clearly showed that the increase in the AGPase small subunit did not match that of the overexpressed large subunit, only reaching around 2-fold of CN (contrasting the 3.5- to 7-fold achieved by the large subunit). Since Zhu et al.’s (2007) model suggested over 9 fold increases in this protein would be necessary, a potential 2-fold increase in active protein might not have been sufficient to relieve limitations on P_i_ recycling. This is further supported by works on Arabidopsis and rice (Obana et al., 2006; Gibson et al., 2011) which showed that plants with 2-fold increases in AGPase activity displayed CO_2_ assimilation rates similar to WT when grown at ambient CO_2_ levels. However, these plants did exhibit an increase in end of day leaf starch which we do not consistently detect in the transgenic tobacco lines. It remains unclear if this is a product of interspecies differences or whether it is due to the nature of the manipulation, as Gibson *et al*. (2011) expressed a deregulated form of AGPase. One might expect the wide range of levels of SBPase protein (1.2 and 6.3-fold of CN) could also explain some of the variation between lines. However, the SBPase amounts in the selected lines SFA28, SFA30, SFA31 and SFA37 were not significantly different between lines at between 3.2 to 3.7-fold of CN levels and closely matching the amount of SBPase required by the model (3.798 fold) (Zhu et al., 2007).

We also would like to highlight the potential for these manipulations to work differently in different species. In Arabidopsis, leaf starch is effectively a short-term carbohydrate reservoir often termed ‘transitory starch’, as the starch accumulated during the day is degraded during the subsequent night, providing a continued supply of carbohydrate in the absence of photosynthesis (Sebastian and Samuel, 2012). Increasing this specific carbohydrate pool then leads to increases in plant growth (Gibson et al., 2011). Contrastingly, tobacco is a known starch accumulator with reserves of this carbohydrate not being fully consumed at night and, in some cultivars, potentially also being sink limited. Since Zhu et al.’s models (Zhu et al., 2007; Zhu et al., 2013) were not based on a specific subset of species, but based on the general dynamics of C3 photosynthesis (taken from published enzyme data from different species grown in different conditions) it might oversee specific aspects of metabolism which would mean the minimal strategy tested here would not work on a starch accumulator species like tobacco, but might still be appropriate for increasing photosynthetic capacity in other species. A recent modelling study reviewing the e-photosynthesis model (Zhu et al., 2007; Zhu et al., 2013) specifically for the crop Solanum tuberosum (potato) also highlights the need for increases to the target enzymes described in this paper -amongst others-(Vijayakumar et al., 2024). Further empirical studies with species with larger sinks, such as potato, will be needed to explore this.

## MATERIALS AND METHODS

### Assembly of binary construct for simultaneous overexpression of SBPase, FBP Aldolase and AGPase

The coding sequences for SBPase (AT3G55800) and AGPase large subunit 1 (AT5G19220) from *A. thaliana*, and FBP Aldolase (NM_001346974) from *Solanum lycopersicum* were synthesized as level 0 modules and used in the generation of the SFA construct via Golden Gate cloning (Engler et al., 2009; Weber et al., 2011; Engler et al., 2014). The transgenes were under the control of CaMV35S and FMV (Richins et al., 1987) constitutive promoters (SBPase and FBP Aldolase respectively) and the *A. thaliana* RbcS2 promoter (AGPase). The construct details are given in Supplementary Figure. 1. This recombinant plasmid was introduced into WT tobacco (*N. tabacum* cv. Samsun) using *A. tumefaciens* strain LBA4404 via leaf-disc transformation (Horsch et al., 1985), and the shoots were regenerated on Murashige Skoog medium containing hygromycin (20 mg l^−1^) and cefotaxime (400 mg l^−1^). Hygromycin-resistant primary transformants (T0 generation) with established root systems were transferred to soil and allowed to self-fertilize. 60 independent lines were generated of which 10 single T-DNA insert lines were taken forward for further analysis in the T2 generation. Control plants (CN) used in this study were null segregant plants from the selected transgenic lines, verified by PCR for non-integration of the transgene and absence of overexpression of all three transgenes by qRT-PCR, immunoblots and proteomics analysis.

### Plant growth conditions and biomass measurements

T2 seeds (homozygous and CN) were germinated on soil in controlled environment chambers at an irradiance of 250 μmol m^−2^ s^−1^, 22 °C air temperature and relative humidity of 60%, in a 16-h photoperiod. 10 days after sowing, seedlings were transferred to individual 8-cm pots and grown under the same conditions for 13–14 days. 23-24 days after sowing, plants were transferred into 4-L pots and moved to a controlled environment glasshouse (16 h photoperiod, 25 °C–33 °C day/20 °C night, with natural light supplemented under low light induced by cloud cover with high-pressure sodium light bulbs, giving 380–1,000 μmol m^−2^ s^−1^ from the pot level to the top of the plant). Plants were distributed in randomized positions in 6 - 12 blocks and watered regularly with Hoagland’s nutrient medium (Hoagland DR, 1950). Plants were positioned such that at maturity, a near-to-closed canopy was achieved. Plants were harvested for biomass at 25, 46 and 56 days after sowing. Stem length and number of leaves per plant were recorded and the total leaf area per plant was measured with a conveyor-belt scanner (LI-3100C Area Meter; LI-COR, Lincoln, NE). Plants were subsequently separated into leaf and stem fractions and dried to constant weight at 70–80 °C, after which dry weight was determined for each fraction.

A smaller-scale experiment to measure root biomass was conducted in controlled environment chambers. Irradiance ranged from 250 μmol m⁻² s⁻¹ at pot level and 450 μmol m⁻² s⁻¹ at the canopy top of plants the day before measurements were taken. Temperature was kept constant at 23°C with a relative humidity of 65-70%. Pots were filled with soil and weighed, ensuring a net weight of 145g at the time seeds were sown. Six plants per line were harvested after 37 days, cut at the stem 1 cm above soil level, and dried at 60-70 °C for 10 days. Weights were recorded for pots containing roots and controls with the same amount of soil but no dried roots.

### Chlorophyll fluorescence imaging

Chlorophyll fluorescence parameters were obtained using a chlorophyll fluorescence imaging system (Technologica, Colchester, UK (Barbagallo et al., 2003; Baker and Rosenqvist, 2004)) fitted with a gas controlled system where the seedlings were exposed to ambient levels of CO_2_ (400-470 ppm), 2% O_2_ and a vapor pressure deficit (VPD) around of 1 kPa. The operating efficiency of PSII photochemistry, *F*_q_ʹ/*F*_m_ʹ, was calculated from measurements of steady-state fluorescence in the light (*F*ʹ) and maximum fluorescence (*F*_m_ʹ) following a saturating 800 ms pulse of 6300 μmol m^−2^ s^−1^ photosynthetic photon flux density (PPFD) and using the following equation: *F*_q_ʹ/*F*_m_ʹ = (*F*_m_ʹ − *F*ʹ)/*F*_m_ʹ (Oxborough and Baker, 1997; Baker et al., 2001). Images of *F*_q_ʹ/*F*_m_ʹ were taken under stable PPFD of 500 μmol m^−2^ s^−1^ or 800 μmol m^− 2^ s^−1^ depending on the experiment. Plants were measured 21-22 days after sowing.

### Leaf Gas Exchange Measurements

Measurements were conducted using a portable infrared gas analyser with an integrated light source (LI-COR 6800; LI-COR, Lincoln, NE). The gas flow rate was kept constant at 500 μmol s^−1^ and the air relative humidity in the leaf chamber was maintained at 55 to 70% depending on the experiment. Plants were measured within the first seven hours of the photoperiod to minimise diurnal effects on stomatal conductance and photosynthetic activity, apart from diurnal measurements. Young developing leaves were measured as well as mature leaves (positions 8 to 14 from the bottom of the plant) before the plants began flowering.

#### *A/C_i_* Response Measurements

To assess the response of CO_2_ assimilation (*A*) and stomatal conductance (*g_s_*) to intercellular CO_2_ concentration (*C_i_*), leaves were initially acclimated at a saturating irradiance of 1500 μmol m^−2^ s^−1^ and a reference CO_2_ concentration of 400 ppm to mimic ambient levels. Following stabilisation, CO_2_ levels were adjusted incrementally (400, 300, 200, 100, 50, 400, 500, 600, 800, 1000, 1200, 1500, and 2000 ppm). Measurements were recorded after *A* and *g_s_* reached a new steady-state (1-2 minutes between collection of data points) with a leaf temperature range of 26-29 °C. From these response curves, maximum Rubisco carboxylation rate (*V_cmax_*), maximum electron transport rate (*J_max_*) and triose phosphate utilisation (TPU) were determined using the equations of Farquhar et al. (1980) and the R package Plantecophys (Duursma, 2015).

#### PPFD Step-Response Measurements

To assess the response of *A* and *g_s_* to changes in light intensity, leaves were initially acclimated at a low PPFD of 100 μmol m^−2^ s^−1^ and a reference CO_2_ concentration of 400 ppm until *A* and *g_s_* reached steady-state levels. After recording *A* and *g_s_* at these levels, the PPFD was increased in a single step to 1500 μmol m^−2^ s^−1^ and *A* and *g_s_* were recorded every 20 seconds for 30 minutes to capture their response during light induction. The leaf temperature ranged from 24-27 °C.

#### Diurnal Measurements

To assess changes in *A* and *g_s_* across the photoperiod, in situ leaf gas exchange measurements were conducted every two hours between 7:30 am and 7:30 pm. PPFD and temperature levels in the leaf chamber were maintained throughout the measurements to match the environmental conditions in the greenhouse.

### Stomatal density measurements

Stomatal density was measured using leaf surface impressions. Silicone impression material (Xantopren, Heraeus, Germany) was used following the methods described by Weyers *et al*. (1985). Following gas exchange measurements, impressions were taken from the same site on four to eight leaves per line. Stomatal density was determined using light microscopy (Olympus BX60, Essex, UK) at 100x magnification. Each impression was analysed using an average of six technical replicates.

### Protein extractions, immunoblotting and proteomic analysis

Leaf discs were ground in dry ice and protein extractions performed as described in Lopez-Calcagno *et al*. (2017) and using the nucleospin RNA/Protein kit (Macherey-Nagel, http://www.mn-net.com/). Protein quantification was performed using the protein quantification Kit from Macherey-Nagel. Samples were loaded on an equal protein basis (5-10 μg per well depending on the experiment), separated using 12% (w/v) SDS-PAGE, transferred to nitrocellulose membranes, and probed using antibodies raised against SBPase (Lefebvre et al., 2005), FBP Aldolase (Simkin et al., 2015), ssAGPase and Actin (AS111739, and AS132640 respectively from Agrisera, via Newmarket Scientific, UK). Proteins of interest were detected using horseradish peroxidase conjugated to the secondary antibody and ECL chemiluminescence detection reagent (Amersham, Buckinghamshire, UK). Membranes were also ponceau stained before blocking to confirm protein transference and loading.

#### Sample preparation for proteomic analysis and LC-MS/MS analysis

Protein pellets were solubilised in 50 mM ammonium bicarbonate, followed by reduction with DTT (5 mM) and alkylation with iodoacetamide (15 mM). The samples were then digested with Sequencing Grade Modified Trypsin (Promega) for 24 hrs. Post-digestion, peptide concentrations were then estimated using the Pierce Quantitative Fluorometric Peptide Assay. The samples were subsequently dried, resuspended in 0.1% formic acid, and normalised to the same concentration ready for LC-MS/MS analysis.

LC-MS/MS analysis was conducted using a Dionex Ultimate 3000 RSLC nanoUPLC (Thermo Fisher Scientific Inc, Waltham, MA, USA) system coupled with a QExactive Orbitrap mass spectrometer (Thermo Fisher Scientific Inc, Waltham, MA, USA). The extracted peptides were initially separated by reverse-phase chromatography at a flow rate of 300 nL/min on a Thermo Scientific reverse-phase nano Easy-spray column (Thermo Scientific PepMap C18, 2µm particle size, 100A pore size, 75µm i.d. x 50cm length). Peptides were loaded onto a pre-column (Thermo Scientific PepMap 100 C18, 5µm particle size, 100A pore size, 300µm i.d. x 5mm length) from the Ultimate 3000 autosampler with 0.1% formic acid for 3 minutes at a flow rate of 15 µl/min. After this period, the column valve was switched to allow elution of peptides from the pre-column onto the analytical column. Solvent A comprised 0.1% formic acid in water, and solvent B consisted of 80% acetonitrile, 20% water and 0.1% formic acid. The linear gradient employed was 2-40% B in 90 minutes, with a total run time of 120 minutes including column washing and re-equilibration.

The eluted peptides were sprayed into the mass spectrometer using an Easy-Spray source (Thermo Fisher Scientific Inc.). All m/z values of eluting ions were measured in an Orbitrap mass analyzer, set at a resolution of 35000, scanning between m/z 380-1500. Data dependent scans (Top 20) were employed to automatically isolate and generate fragment ions by higher energy collisional dissociation (HCD, NCE:25%) in the HCD collision cell, with the resulting fragment ions measured in the Orbitrap analyzer at a resolution of 17500. Singly charged ions and ions with unassigned charge states were excluded from MS/MS selection, and a dynamic exclusion window of 20 seconds was employed.

All MS/MS samples were analysed using Mascot (Matrix Science, London, UK; version Mascot in Proteome Discoverer 2.4.1.15). Mascot was set up to search against *Nicotiana tabacum* proteome (Uniprot, downloaded on 02/06/2021) and a common contaminant database, assuming trypsin digestion. Mascot searches were performed with a fragment ion mass tolerance of 0.100 Da and a parent ion tolerance of 20 PPM. Carbamidomethyl of cysteine was specified as a fixed modification, while deamidated of asparagine and glutamine and oxidation of methionine were specified as variable modifications. Peak areas of identified peptides were calculated for label-free quantification.

Scaffold (v4.10.0, Proteome Software Inc., Portland, OR) was used to validate MS/MS based peptide and protein identifications. Peptide identifications were accepted if they could be established at greater than 95.0% probability by the Scaffold Local FDR algorithm. Protein identifications were accepted if they could be established at greater than 99.0% probability and contained at least 2 identified peptides. Protein probabilities were assigned by the Protein Prophet algorithm (Nesvizhskii et al., 2003). Proteins containing similar peptides and could not be differentiated based on MS/MS analysis alone were grouped to satisfy the principles of parsimony. Proteins sharing significant peptide evidence were grouped into clusters.

### RNA extraction and transcriptomics

Leaf discs of the relevant transgenic and azygous lines were ground in dry ice and RNA extractions were performed following the nucleospin RNA/Protein kit (Macherey-Nagel). RNAseq was performed by Novogene. After checking RNA read quality using FastQC (https://www.bioinformatics.babraham.ac.uk/projects/fastqc/), a classification-based quantification was performed using kallisto (Bray et al., 2016). In short, a kallisto index was built with the reference transcriptome of *Nicotiana tabacum* (Edwards et al., 2017) appending the sequences of the introduced genes. Kallisto quant was used to quantify abundance of pair-end reads with default parameters. The resulting abundance files were used to generate a count table in R using package tximport. Genes with low counts (less than 10 reads) were removed and differential expression analysis was performed in R using DESeq2. Genes with an adjusted p.value < 0.001 and a Fold2logChange Ratio > 1.5 (or < −1.5) were highlighted as differentially expressed and included in Supplemental Table 1.

### Soluble sugar and starch determination

Samples from fully expanded leaves were taken within a 1h window at the end of the day and night periods and flash frozen. Ethanolic extracts from 20 mg of frozen plant material were used to determine sucrose, glucose, fructose and starch as in Cross *et al*. (2006). For starch determination, pellets were solubilised by heating them to 80°C in 0.1M NaoH for 40 min. After neutralising the pH with an HCl/sodium-acetate solution, the suspension was digested overnight witn amyloglucosidase and amylase. The glucose content of the supernatant was then used to determine the starch content in the sample.

### Statistical analysis

All statistical analyses were performed using R (https://www.r-project.org/, version 4.0.5). Analysis of variance considering block effect when relevant, followed by post hoc T-test or Tukey tests were carried out.

## ACKNOWLEDGEMENTS

We thank the transformation team at the University of Illinois for generating the transgenic T0 lines and Phillip A. Davey (University of Essex, UK) for help with gas exchange and Chlorophyll fluorescence at low O_2_. This study was funded by a subcontract to the University of Illinois as part of the Bill and Melinda Gates Foundation RIPE (Realizing Increased Photosynthetic Efficiency) programme.

## AUTHOR CONTRIBUTIONS

PELC generated constructs and did initial line screening. PELC & PL led the experimental design and physiological measurements with support from SW and BM. HG conducted the proteomic analysis and CA the RNAseq. BM conducted the carbohydrate quantification. Each co-author carried out data analysis on their respective contributions. PELC and CAR wrote the article with input from TL. CAR secured the funding.

